# EAP: a versatile cloud-based platform for comprehensive and interactive analysis of large-scale ChIP/ATAC-seq data sets

**DOI:** 10.1101/2024.03.31.587470

**Authors:** Haojie Chen, Tao Huang, Zhijie Guo, Anqin Zheng, Weiran Chen, Liangxiao Ma, Shiqi Tu, Guangyong Zheng, Yixue Li, Zhen Shao

## Abstract

Epigenome profiling techniques such as ChIP-seq and ATAC-seq have revolutionized our understanding of gene expression regulation in tissue development and disease progression. However, increasing amount of ChIP/ATAC-seq data poses challenges in computational resources, and the absence of systematic tools for epigenomic analysis underscores the necessity for an efficient analysis platform. To address these issues, we developed the Epigenomic Analysis Platform (EAP, https://www.biosino.org/epigenetics), a scalable cloud-based tool that efficiently analyzes large-scale ChIP/ATAC-seq data sets. EAP employs advanced computational algorithms to derive biologically meaningful insights from heterogeneous datasets and automatically generates publication-ready figures and tabular results, enabling comprehensive epigenomic analysis and data mining in areas like cancer subtyping and therapeutic target discovery.

## Introductions

Epigenome profiling plays a crucial role in investigating gene expression regulation in tissue development and disease progression^1,2^. Advanced deep sequencing techniques, including ChIP-seq and ATAC-seq, have enabled us to dissect the epigenetic heterogeneity of developmental cells and disease cohorts by identifying epigenomic alterations and the potential associated transcription factors (TFs)^3,4^. These high-throughput techniques provide valuable insights into the mechanisms that control gene expression and disease progression^5,6^.

In recent years, large consortium projects such as ENCODE^7^ and TCGA^1^ as well as many cancer genomics studies have generated a tremendous amount of ChIP/ATAC-seq data in various normal and cancer cells/tissues, providing opportunities for data mining and a broader understanding of epigenetic regulation of gene expression. However, as the scale of data generation continues to grow, researchers require more computational resources to analyze these large datasets. Additionally, there is a lack of systematic epigenomic analysis tools to explore the huge amount of ChIP/ATAC-seq data deposited in NCBI GEO^8^ and CNCB GSA^9^. Despite the availability of a large number of computational tools and methods for ChIP/ATAC-seq data analysis, it remains challenging for experimental biologist to deploy and integrate these tools into workable pipeline, particularly in heterogeneous cohort studies (e.g., those involving patients that may be in different disease states or subtypes) where conventional analysis tools are often inadequate.

To address these gaps, we have developed EAP (Epigenomic Analysis Platform, https://www.biosino.org/epigenetics), a scalable, customizable and interactive ChIP/ATAC-seq data analysis platform based on cloud technology. EAP uses state-of-the-art computational and statistical algorithms to transform ChIP/ATAC-seq data generated from a large panel of samples into biologically meaningful and interpretable results^10–12^. Currently, EAP provides a series of data analytical tools with an interactive web interface to facilitate its use by researchers with limited programming experience, encompassing data preprocessing, supervised differential analysis, differential TF motifs enrichment analysis, differential TF activity analysis, unsupervised hypervariable analysis, clustering analysis, signature genes scoring analysis, etc. All of these analytical tools are specifically developed for modeling and understanding the epigenetic variations among a large number of patient samples or cellular states. The comprehensive epigenomic analysis through EAP can greatly facilitate data mining in different research areas, such as cancer subtyping and therapeutic target discovery.

## Results

### Overview of EAP architecture and the analysis modules

EAP is primarily tailored for cancer patient cohort studies, necessitating a login for the upload and analysis of sensitive personal information (Supplementary Figure 1a). In order to utilize EAP, users must register an account and request storage space by submitting an application form to the administrator. The input files required for EAP include raw sequencing data in FASTQ format and a metadata file in CSV format, containing details on study design and sample phenotypes. Due to the typically large size of these files, we have developed a dedicated data transfer client tool that supports break-point resumable transfers (Supplementary Figure 1b). Additionally, a md5sum checking procedure has been integrated to ensure the integrity of uploaded files (Supplementary Figure 1c). Currently, EAP offers two analytical modules to transform ChIP/ATAC-seq data from heterogeneous samples into biologically meaningful and interpretable results. EAP leverages private cloud computing technology^13^ to establish an analytical framework for processing, interpreting, and visualizing large datasets, ensuring ample computing power for these tasks. Additionally, Docker container technology^14^ is utilized to create automated analysis pipelines and tools, ensuring reproducibility of results and streamlining end-to-end bioinformatics analysis for large epigenomic datasets. The cloud computing architecture supports two analytical modules for both basic and advanced epigenomic data analysis in EAP (Figure 1a).

**Figure 1.**
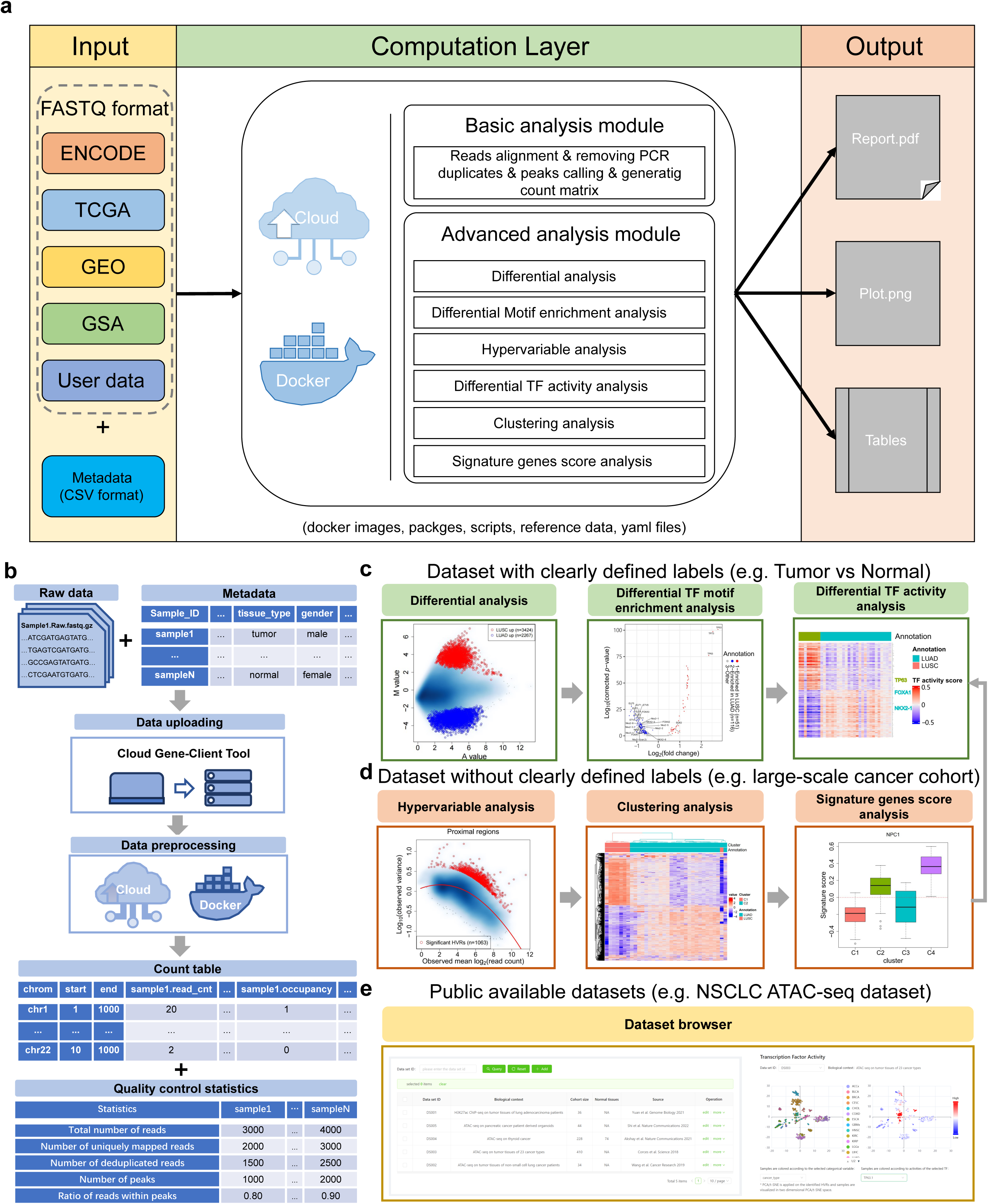
Overview of EAP architecture and the various analysis modules. **(a)** Inputs to EAP include raw sequencing data (in FASTQ format) and metadata (describes study design and sample phenotypes, in CSV format). EAP consists of Basic Analysis module and Advanced Analysis module. Basic Analysis module performs quality control, read mapping, peaks calling, creates a summary report for filtering poor quality samples and generates analysis-ready count tables, which are then used as inputs for Advanced Analysis module. A comprehensive collection of ChIP/ATAC-seq data analysis tools are encapsulated in the Advanced Analysis module. Each analysis tools will produce publication-ready results (figures and tables). (**b)** Workflow of Basic Analysis module, requires the upload of raw sequencing data and metadata using the Cloud Gene-Client tool. Upon completion of this analysis, analysis- ready count tables and a summary quality control report are generated for downstream analysis. **(c-d)** Two distinct research scenarios are illustrated: (c) For labeled data sets, differential analysis can be employed to identify differential signals among samples with different labels. Subsequently, differential TF motif enrichment analysis and differential TF activity analysis can be conducted to investigate the TFs linked to differential binding or open chromatin sites. (d) In datasets lacking pre-defined sample labels, hypervariable analysis can be utilized to identify hypervariable ChIP/ATAC-seq signals across the samples. These signals can then be leveraged for clustering analysis and cluster samples into distinct groups, and signature genes scoring analysis is utilized to characterize these clusters based on established gene sets. Supervised analysis tools can also be utilized to identify binding/open chromatin sites or transcriptional regulators specific to each cluster. **(e)** EAP offers a Data Set Browser that provides an interactive interface for convenient visualization of TF activity scores in each dataset.

### EAP enables efficient and comprehensive large-scale ChIP/ATAC-seq data analysis

The Basic Analysis module is designed and implemented as an automatic pipeline tailored for routine data processing demands, which includes quality control, read alignment, peak calling and read counting. Upon completion of the pipeline, an analysis report (in PDF format), including quality control plots and summary statistic values, will be presented for further study (Figure 1b). NGS data preprocessing is a time-consuming step that demands substantial computational resources. EAP leverages Cloud computing technology to handle large ChIP/ATAC-seq datasets, which can comprise tens or even hundreds of ChIP/ATAC-seq profiles from different samples. Table 1 demonstrates that EAP’s basic analysis module can efficiently process and analyze large datasets from various studies.

**Table 1.** The time cost of data preprocessing using EAP’s Basic Analysis module on public data sets with various sample and file sizes.

| data set | running time | sample size | total file size | source |
| --- | --- | --- | --- | --- |
| GBM ATAC-seq | 61h45min08s | 60 | 630.6GB | Lu et al.<sup>15</sup> |
| LUAD H3K27ac ChIP-seq | 17h58min36s | 42 | 790.3GB | Yuan et al.<sup>16</sup> |
| PDAC organoid ATAC-seq | 29h19min35s | 44 | 722.6GB | Shi et al.<sup>17</sup> |
| Thyroid cancer ATAC-seq | 120h10min25s | 227 | 3.9TB | Sanghi et al.<sup>18</sup> |

GBM: Glioblastoma; LUAD: lung adenocarcinoma; PDAC: Pancreatic ductal adenocarcinoma.

For the Advanced Analysis Module, it encapsulates a comprehensive collection of analytical tools meeting needs of customized ChIP/ATAC-seq data analyses (Figure 1c-d). In practice, users have the flexibility to perform both routine and customized analyses on their data seamlessly, as the input read count table for these advanced analyses is derived from the Basic Analysis module. Furthermore, EAP offers a user-friendly interface that enables users to configure parameters for the advanced analyses, including the number of principal components utilized for sample clustering, the design of the differential analyses, and the specifications for *p*-value/FDR cutoffs, among others (Supplementary Figure 2). Users can choose the specific combination of analytical tools that best suit their research scenarios. For instance, for ChIP/ATAC-seq datasets with clearly defined sample labels, differential analysis can be used to detect the differential signals between samples with different labels, followed by differential TF motif enrichment analysis or differential TF activity analysis to further explore the TFs associated with the differential genome binding or open chromatin sites. This scenario is referred to as supervised analysis (Figure 1c). In contrast, for datasets with no pre- defined sample labels (e.g., those generated from patients with the same disease state/type) or highly sophisticated sample labels (e.g., those covering tens of different cellular states or disease types), users can apply hypervariable analysis to identify hypervariable ChIP/ATAC-seq signals across the samples, which could be then used as features for clustering analysis to dissect the underlying heterogeneity structure among the samples. Then, samples are grouped into different clusters, coupled with signature genes scoring analysis to annotate those clusters based on established gene sets (Figure 1d). Supervised analysis tools could also be applied to the resulting sample clusters to detect binding/open chromatin sites specific to each cluster (Figure 1c-d).

### EAP provides interactive data set browser

EAP has been successfully applied to analyze ChIP/ATAC-seq data from many cancer epigenomic studies, including H3K27ac ChIP-seq on LUAD cohort^16^, ATAC-seq on NSCLC cohort^19^, ATAC-seq on TCGA pan-cancer cohort^1^, ATAC-seq on thyroid cancer cohort^18^ and ATAC-seq on pancreatic cancer patient derived organoids^17^. These processed data sets are available on the Data Set Browser in EAP, and EAP offers an interactive interface for easy visualization of TF activity scores in each data set. This module allows users to investigate the role of interest of transcriptional regulators in oncogenesis by choosing an appropriate data set (Figure 1e).

### Comparison of comprehensiveness of supported ChIP/ATAC-seq features across EAP and other existing analysis platforms/tools

While numerous ChIP/ATAC-seq analysis platforms/tools exist, EAP is designed to provides more flexible and comprehensive down-stream analyses, enabling users to delve deeper into their data and extract biological meaningful insights. Additionally, EAP allows for adding new features in future releases (Table 2). One of the key advantages of EAP is its exceptional ability to handle large-scale ChIP/ATAC-seq datasets ranging from gigabytes to terabytes in size. This capability sets EAP apart from other tools, as it enables users to efficiently analyze and process vast amounts of data.

**Table 2.** Comparison of comprehensiveness of supported ChIP/ATAC-seq features from currently available analysis platforms/tools.

|  |  | ChIPseek<sup>20</sup> | PEPATAC<sup>21</sup> | Cistrome<sup>22</sup> | CisGenome<sup>23</sup> | EAP |
| --- | --- | --- | --- | --- | --- | --- |
| Data preprocessing | trimming adapter | × | ✓ | ✓ | ✓ | ✓ |
|  | read alignment | × | ✓ | ✓ | ✓ | ✓ |
|  | peaks calling | × | ✓ | ✓ | ✓ | ✓ |
|  | peaks annotation | ✓ | ✓ | × | ✓ | ✓ |
|  | read distribution | × | ✓ | ✓ | ✓ | ✓ |
|  | read counting | × | × | ✓ | × | ✓ |
| Downstream analysis | motif enrichment analysis | ✓ | ✓ | ✓ | × | ✓ |
|  | differential analysis | × | × | ✓ | × | ✓ |
|  | differential TF motif enrichment analysis | × | × | × | × | ✓ |
|  | hypervariable analysis | × | × | × | × | ✓ |
|  | differential TF activity analysis | × | × | × | × | ✓ |
|  | clustering analysis | × | × | ✓ | × | ✓ |
|  | signature genes score analysis | × | × | × | × | ✓ |
|  | allow for adding new features | × | × | ✓ | × | ✓ |
| Other | web based analysis tools | ✓ | × | ✓ | × | ✓ |
|  | genome browser visualization | ✓ | ✓ | ✓ | ✓ | × |

### Case Study 1: ATAC-seq profiles of different tissue types from thyroid cancer patients

To demonstrate the power of EAP with labeled datasets, we collected an ATAC-seq dataset of thyroid cancer^18^, comprising 70 primary tumor tissues, 70 patient-match normal tissues and 83 metastatic cancer tissues. We utilized EAP to preprocess the sequencing data and then conducted pairwise differential analysis of different tissue types using count table acquired from the Basic Analysis module. Our analysis revealed that in comparison to the alterations in chromatin accessibility between primary tumor tissues and metastatic cancer tissues, a notably higher number of differentially accessible sites were identified when contrasting primary tumor tissues/metastatic cancer tissues with normal tissues (Figure 2a). We further performed differential TF motif enrichment analysis to detect TF motifs that are differentially enriched in one set of peaks regions relative to another set (e.g., tumor tissue up-regulated peaks vs normal tissue up-regulated peaks). We found that normal thyroid development transcription factors FOXE1 and HHEX were significantly enriched in normal tissue up-regulated sites^24^. However, the potential binding sites for transcription factors of the AP-1 family are more enriched in the up- regulated sites in primary tumors tissue/metastatic cancer tissue, such as FOS, JUN, FOSL2, JDP2, and BATF (Figure 2b)^25^.

**Figure 2.**
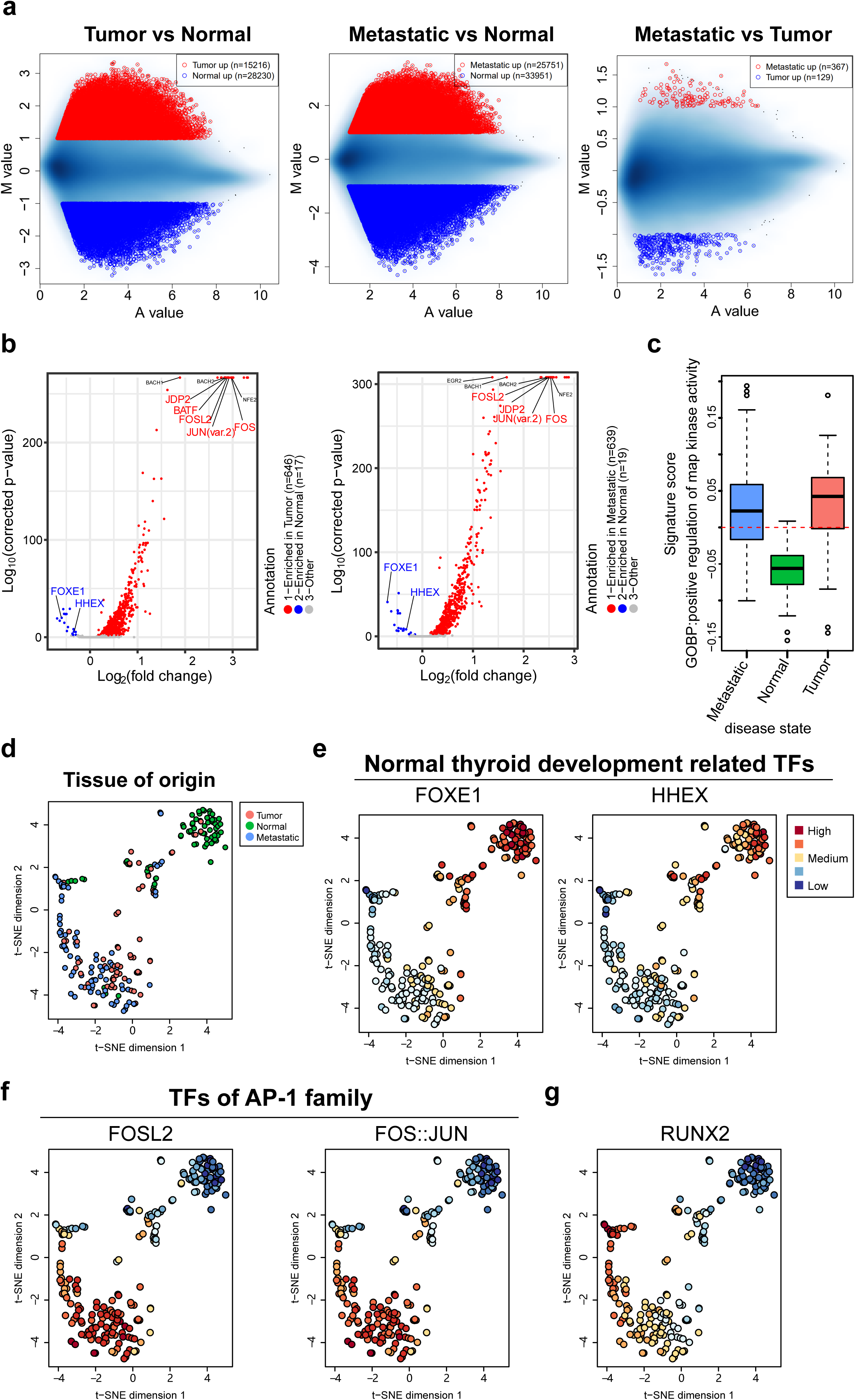
Illustration of application of EAP on labeled data sets. **(a)** MA plots highlight the differentially accessible sites when comparing ATAC- seq profiles across normal tissues, tumor tissues, and metastatic tissues. **(b)** Volcano plots show the differential TF motif enrichment analysis results of differentially accessible sites between normal tissues and tumor/metastatic tissues. **(c)** Boxplot shows the MAPK pathway activities across different tissue types. **(d)** Two dimensional t-SNE visualization of ATAC-seq samples using hypervariable regions identified by EAP, samples are colored by tissue of origin. **(e-g)** Two dimensional t-SNE visualization of different ATAC-seq samples colored by TF activity scores calculated by EAP.

The AP-1 family is the main target proteins of MAPKs^26^, and previous studies have shown that thyroid cancer is driven by MAPK signaling pathway^27^. Therefore, these transcription factors enriched in up-regulated sites in primary tumors tissue/metastatic cancer tissue may be related to the activation of MAPK signaling pathway in thyroid cancer. To verify the activity of MAPK signaling pathway in different tissue types, signature genes activity score analysis tool implemented in EAP was employed to assign the MAPK signaling pathway activity estimates to individual samples. Our results indicated that primary tumor tissues/metastatic cancer tissues have indeed a higher MAPK signaling pathway activity than normal tissues (Figure 2c). Additionally, we performed differential TF activity analysis to identify tissue type specific transcriptional regulators. Consistent with previous findings, TF activity scores of normal thyroid development related transcription factors are higher in normal tissues, while TF activity scores of AP-1 family TFs are increasing in primary tumors tissues/metastatic cancer tissues (Figure 2d-f). It is worth noting that the activity scores of RUNX2 are progressively increasing from patient-match normal tissues to primary tumors tissues, and at last metastatic cancer tissues, which exhibited significantly higher activity (*P*=1.98E-20) (Figure 2g). Enhanced regulation activity of RUNX2 have been reported to be functionally linked to tumor invasion and metastasis in thyroid cancer^28^. Overall, using the combination of EAP differential analysis tools, we found that increasing of AP- 1 family TFs regulation ability, up-regulation of MAPK signaling pathway and increased activity of RUNX2 are related to the carcinogenesis and progression of thyroid cancer. These findings demonstrate the functional profiling capacity of EAP in capturing epigenetic alterations and the associated transcription factors in labeled data set.

### Case Study 2: ATAC-seq profiles of patient-derived pancreatic cancer organoids

However, in most cancer studies, due to the heterogeneity among patients and the complexity or lack of clinical pathological information, it is difficult to use existing information as sample labels for down-stream analysis. Consequently, EAP provide an unsupervised analysis strategy for analyzing label-free dataset. We applied EAP to reanalyze an ATAC-seq dataset containing 44 patient-derived pancreatic cancer organoids (Figure 3a)^17^. Four different pancreatic ductal adenocarcinoma (PDAC) subgroups were identified based on previously defined signature gene sets^29,30^, including classical progenitor like subgroups (C1), basal-like subgroups (C2), classical immunogenic like subgroups (C3) and aberrantly differentiated endocrine and exocrine (ADEX) like subgroups (C4 and C5) (Figure 3b-c). According to basal subtype signature gene scores, we divided the four subgroups into basal-like and non-basal samples, basal-like PDAC showed significantly worse prognosis than those in non-basal patients (*P*=0.045), which is in line with previous studies (Figure 3d)^5,31^. To further elucidate differences in transcriptional regulators between different subgroups, we identified subgroup-specific transcription factors using the differential TF activity analysis tool. GATA6, HNF1A, and HNF1B, which are critical for normal pancreatic development and differentiation, were enriched in classical progenitor like subgroups^32–34^. In the classical immunogenic-like subgroup, HNF4A and HNF4G were identified, and these transcription factors were reported not only to be associated with the classical pancreatic cancer subtype, but also to play an important role in the invasion of pancreatic cancer^35^. RUNX1, RUNX2 and RUNX3 were significantly enriched in the basal-like subgroup, and the high expression of these transcription factors have been reported to promote pancreatic cancer cell migration, invasion and drug resistance, which can be used as a potential target for pancreatic cancer treatment^36,37^. Transcription factors related to pancreatic endocrine or neuroendocrine, such as NEUROD2 and ASCL1^38^, were significantly enriched in the ADEX like subgroup. In addition, Rbpj1 is critical in the differentiation of pancreatic acinar cells and it was also enriched in ADEX like subgroup(Figure 3e)^39^. These results indicate that both endocrine and exocrine transcription factors are highly activated in this subgroup, rather than one of the endocrine or exocrine-related transcription factors, which is common in normal pancreas development. Overall, these findings exemplified the utility of EAP in revealing both cancer epigenetic subtypes and subtype specific transcriptional regulators in a label-free data set.

**Figure 3.**
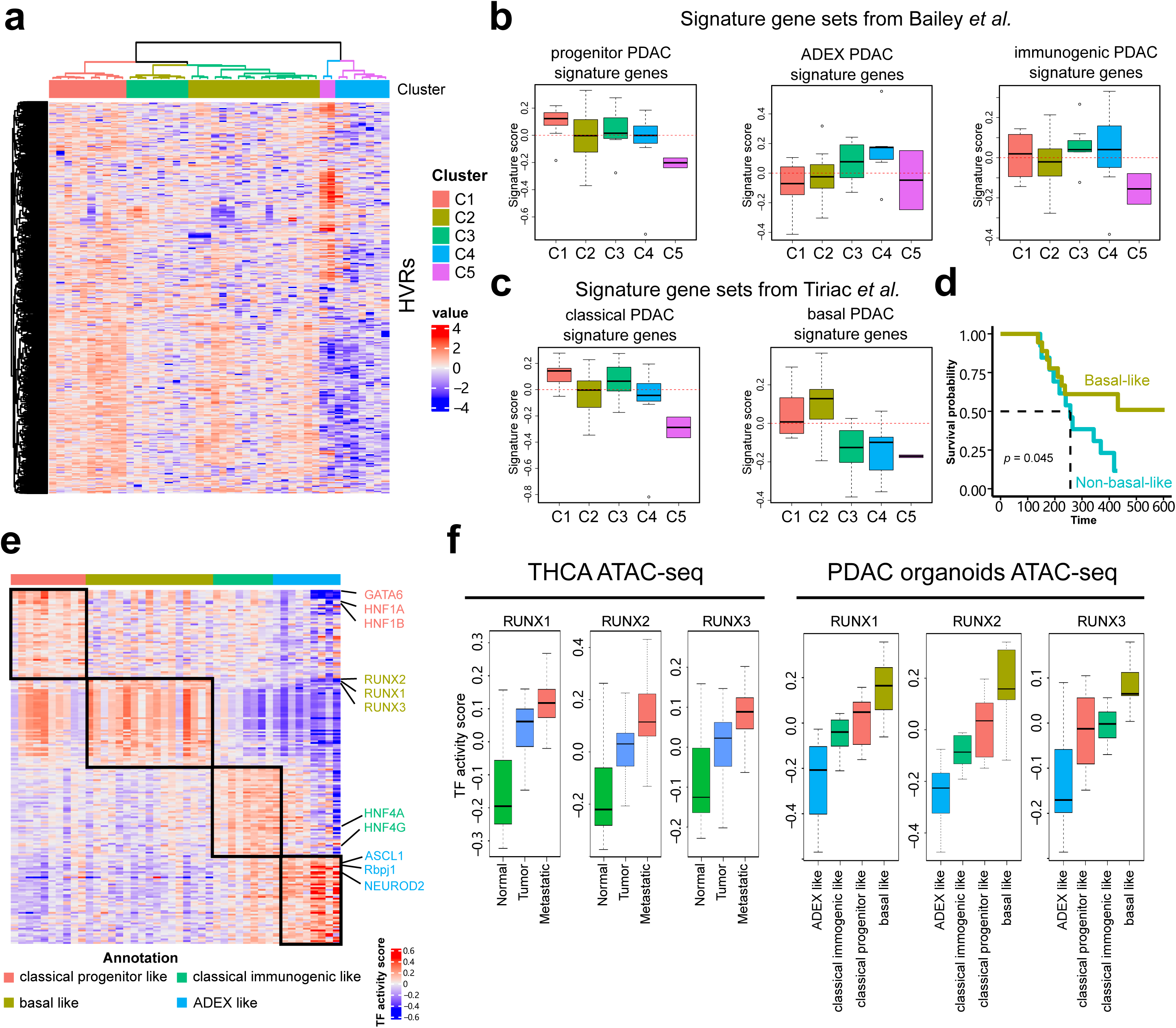
Illustration of application of EAP on label-free data sets. **(a)** Heatmap and dendrogram showing clustering of samples with similar chromatin accessibility profiles and clustering of peak regions with similar expression patterns. **(b-c)** Boxplots showing the expression activity of signature gene sets curated by previous studies. **(d)** Kaplan-Meier curves displaying differences in progression-free survival (PFS) between patients whose organoids defined as Basal-like or Non-basal-like based on ATAC-seq profiles. TLSs&LAs signatures score. **(e)** Heatmap of TF activity scores of top-ranked cluster specific TFs. **(f)** Boxplots showing the TF activity scores across samples in two public cancer related data sets, samples are grouped by tissue of origins (left panel) or subtypes defined EAP (right panel).

## Discussions

In summary, EAP presents a revolutionary data analysis platform to support large-size epigenomic dataset analyses based on the cloud combining technology. We demonstrated the practicability and high-efficiency of EAP in dissecting cancer epigenetic heterogeneity, which can be used not only for mining key transcriptional regulators of different cancer pathological subtypes, but also for cancer epigenetic subtyping, and further revealing subtype associated transcription factors.

Furthermore, EAP offers interactive data analysis, allowing end users to iteratively modify analytical parameters to enhance the identification of biologically significant results. This feature is particularly crucial for cancer epigenomic data analysis, where a standardized analytical pipeline may not be suitable or preferred.

EAP has been successfully applied to analyze ChIP/ATAC-seq data from many cancer epigenomic studies, which comprise tens or even hundreds of ChIP/ATAC-seq profiles from different individuals. The processed data sets are available on the Data Set Browser in EAP, and EAP offers an interactive interface for easy visualization of TF activity scores in each data set. It is worth noting that as we applied this module on datasets from multiple cancer types, RUNX family TFs exhibited higher regulation activity in both progressive subtypes and metastatic tissues (Figure 3f and Supplementary figure 3), which indicate RUNX family TFs might as candidate multi-cancer progression related regulators and may be a potential target for cancer therapy.

We anticipate that EAP can be used for more large-scale cancer epigenomic data analyses, leading to more interesting and meaningful biological or clinical discoveries, which broaden our understanding of cancer carcinogenesis and progression related epigenetic machinery.

## Methods

### Implementation

The EAP web server runs on a private cloud system equipped with 5,654 GB of memory, 824 CPUs, and 10 TB of storage space. Docker technology was utilized to create images containing necessary software and packages for ChIP/ATAC-seq data analysis, such as *FastQC* (v0.11.9) for read quality control, *Bowtie* (v1.2.0) for read alignment, and *MACS* (v1.4.2) for peak calling. Analysis workflows, including data preprocessing pipelines and downstream analysis tools, were defined in YAML files specifying configuration parameters and commands to be executed. Subsequently, graphical user interfaces (GUIs) for the data preprocessing pipeline and downstream analysis tools were deployed on the web server. The user-friendly GUI was designed to allow non-technical users to customize analysis parameters (e.g., adjusted *p*-value cutoff, number of clusters, variables of interest for differential analysis) and conduct various ChIP/ATAC-seq data analyses.

EAP implements login authentication as a crucial measure for data security and storage. Users are required to register an account (https://www.biosino.org/epigenetics/#/user/register) and request storage space by submitting an application form to the administrator (application form was available at https://github.com/haojiechen94/EAP/blob/main/doc/File_1_Storage_space_a <u>pplication_form.xlsx</u>). Once users have an account and allocated storage space, they can upload raw sequencing data to their designated storage area and utilize the analysis tools integrated within EAP. Instructions and help menus are readily accessible on the Home page to guide users through the platform (help manus was also available at https://github.com/haojiechen94/EAP/blob/main/doc/Help%20document.pdf). Additionally, we also provide video tutorials that visually demonstrate the processes and make it easier for users to learn and apply the procedures effectively (Supplementary movie 1, Supplementary movie 2 and Supplementary movie 3, https://github.com/haojiechen94/EAP/tree/main/video_tutorials). Given the typically large size of uploaded files, a data transfer client was developed to support break-point resumable transfers. Users are advised to utilize this client for uploading raw sequencing data to their allocated storage space within EAP.

### Basic Analysis module: a standardized ChIP/ATAC-seq data preprocessing pipeline

### Input

The input for the data analysis process includes raw sequencing data in FASTQ format and metadata containing descriptions of the raw sequencing data and phenotypic information for each sample. A metadata template can be accessed on the Home page of the EAP platform (metadata template was also available at https://github.com/haojiechen94/EAP/blob/main/doc/File_2_metadata.csv).

### Data Processing

Firstly, *FastQC* (v0.11.9) is utilized for quality control of read bases and *Trim- galore* (v0.6.7) is employed for trimming adapters and removing low-quality bases from reads. ChIP/ATAC-seq reads are aligned to the user-specified reference genome, which can be downloaded from the UCSC genome browser (https://hgdownload.soe.ucsc.edu/downloads.html), using *Bowtie* (v1.2.0). Then, PCR duplicates are removed based on genomic position and remaining reads are used for peak calling with *MACS* (v1.4.2). Finally, a count table is generated using *MAnorm2-utils* (v1.0.0).

### Output

The analysis generates three count tables where genomic regions are represented in rows and samples in columns.

- One count table (NA_profile_bins.xls) is prepared for differential analysis.

- Two count tables (proximal_peak_regions_2000bp.txt and distal_peak_regions_2000bp.txt) are prepared for hypervariable analysis.

- A summary report is produced, encompassing quality control assessments for bases, reads mapping, and peaks calling, as well as genomic annotations of peaks and motif enrichment within peaks.

### Advanced Analysis module: a comprehensive collection of ChIP/ATAC-seq data analysis tools

#### Differential Analysis

##### Input

The input for the differential analysis module includes count tables (NA_profile_bins.xls) generated from the output of the Data Preprocessing module and metadata.

##### Data Processing

Raw count tables are normalized using *MAnorm2* to account for differences in library size and correct MA trended bias. Then differential analysis is conducted based on the user-specified variable of interest.

##### Output

The output of the analysis includes:

- A table presenting the results of the differential analysis.

- MVC plot illustrating the global mean-variance trend.

- MA plot highlighting significantly differentially enriched or accessible peaks.

### Differential TF Motif Enrichment Analysis

#### Input

The input for the differential TF motif enrichment analysis module consists of the result table from the previous step of the differential analysis.

#### Data Processing

A user-selected adjusted *p*-value cutoff is applied to define significantly differentially enriched or accessible peaks (DEPs or DAPs). Then *motifscanR* tool is utilized to scan for transcription factor (TF) motif occurrences in these DEPs/DAPs. Finally, Fisher’s exact test is employed to determine if a TF motif is significantly enriched in one set of peak regions relative to another set.

#### Output

The output of the analysis includes:

- A table presenting the results of the differential motif enrichment analysis.

- Enriched/depleted *p*-values indicating the extent to which a TF motif is over/under-represented in a peak set relative to another peak set.

- A volcano plot highlighting significantly differentially enriched TF motifs.

### Hypervariable Analysis

#### Input

The input for the hypervariable analysis module includes count tables (proximal_peak_regions_2000bp.txt and distal_peak_regions_2000bp.txt) obtained from the output of the Basic Analysis module and metadata.

#### Data Processing

MA normalization is performed in pseudo-reference mode. *HyperChIP* is applied to evaluate the significance of signal variability. Peak regions with an adjusted *p*-value below 0.001 (default) are identified as hypervariable regions. Then [rincipal component analysis (PCA) is conducted based on signals within these variable regions and samples are visualized in two-dimensional PCA space or *t*-SNE space.

#### Output

The output of the hypervariable analysis tool includes:

- Hypervariable analysis results exported in both TXT and RData formats.

- MVC plots illustrating the global mean-variance trend in proximal and distal regions, with significant hypervariable regions marked in red.

- Scatter plots displaying the PCA results or t-SNE dimension reduction results based on signals in hypervariable regions.

### Clustering Analysis

#### Input

The input for the clustering analysis includes hypervariable analysis results in RData format and metadata.

#### Data Processing

Principal component analysis (PCA) is conducted on signals within hypervariable regions and the top-ranked principal components (PCs) are utilized to calculate the Euclidean distance between each pair of samples. Then hierarchical clustering is applied to group samples into clusters based on the similarity of their signal patterns in hypervariable regions (HVRs).

#### Output

The output of the clustering analysis tool includes:

- A new metadata containing the clustering results, which can be utilized for further analyses.

- A cluster heatmap displaying the hierarchical clustering result, providing a visual representation of sample similarities and clustering patterns.

### Signature genes score analysis

#### Input

Hypervariable analysis result of proximal regions in RData format, genes to proximal regions links and a list of genes of interest in GMT format.

#### Data processing

Given a gene set of interest, it is usually more desirable to summarized the expression level of that gene set using a single integrated score. This tool standardizes the ChIP/ATAC-seq signals in the proximal regions within a given

dataset by *z*-score transformation. Then summarizes resulting scores of those proximal regions linked to the genes of interest, minus the mean of z-scores of all proximal regions as negative control.

#### Output

The output of the Signature genes score analysis tool includes:

- A boxplot shows the distribution of signature scores among different clusters. This analysis provides visualization to improve interpretation of the clustering results. For example, users can annotate the cluster based on these signature genes scores.

### Differential TF Activity Analysis

#### Input

Hypervariable analysis results in RData format and metadata.

#### Data processing

Peak regions with adjusted *p*-value below 0.001 (default) were defined as hypervariable regions. *motifscanR* was then utilized to conduct motif scanning on these genomic regions, aggregating TF motif-associated signals in each sample to generate a score representing TF regulatory activity. Following this, a *t*-test was employed to identify TFs associated with the user-specified variable of interest.

#### Output

The outputs in this analysis including a table of TF activities in each sample, a table of *t*-statistic of the association test and plots for dimension reduction visualization of samples and the activities of user specified TFs.

#### Evaluation

We showcase application of EAP with two distinct analysis scenarios: (i) In data set with clearly defined labels. In this analysis scenarios, we reanalyzed an ATAC-seq data set of thyroid cancer, which included 70 primary tumor tissues, 70 patient-match normal tissues and 83 metastatic cancer tissues. Using EAP,

we investigated chromatin accessibility alterations during cancer progression and identified potential associated transcriptional regulators. (ii) In data sets without pre-defined labels. For this case study, we obtained ATAC-seq data on

44 patient-derived pancreatic cancer organoids. First, we perform sample clustering based on chromatin accessibility heterogeneity and define subtypes using established signature gene sets. Subsequently, we identified key regulators specific to each subtype.

#### Data availability

This study made use of multiple publicly available datasets. For Case Study 1, the thyroid cancer ATAC-seq was obtained from GEO, under accession number GSE162515. For Case Study 2, the PDAC organoid ATAC-seq data was obtained from GSA, under accession number HRA002013. The LUAD H3K27ac ChIP-seq data was downloaded from EGA, under accession number EGAD00001007066. The GBM ATAC-seq data was obtained from GEO, under accession number GSE163853. The NSCLC ATAC-seq data was obtained from West China Hospital (https://pms.cd120.com/download.html). The pan-cancer ATAC-seq data set was obtained from TCGA GDC (https://gdc.cancer.gov/about-data/publications/ATACseq-AWG).

## Code availability

Extensive documentation and a full user manual are available at https://github.com/haojiechen94/EAP/tree/main/doc. The software is open source, and all code can be found on GitHub at https://github.com/haojiechen94/EAP/tree/main/source_codes.

## Author’s contributions

Z.S., Y.L., G.Z. and S.T. conceived the idea and designed the analysis platform. H.C., T.H., Z.G. and A.Z. developed the web server and analyzed the real datasets. H.C., Z.G., Z.S. and G.Z. wrote the manuscript. H.C. wrote the

documentation with help from Z.G., W.C. and L.M.. All authors read and approved the final manuscript.

## Supporting information

Supplementary movie 1 Data transferring and integrity checking

Supplementary movie 2 Basic Analysis module demo using paired end ChIP seq data

Supplementary movie 3 Advanced Analysis module demo differential analysis

## Supplementary figures

**Supplementary figure 1 The login page and data transfer tools of EAP.**

**(a)** The login page of EAP (https://www.biosino.org/epigenetics/#/user/login), a demonstration account is available by click on "demo" in this page. **(b)** The data transfer client for utilization of EAP, which supports break-point resumable transfers. **(c)** A md5sum checking procedure has been integrated to ensure the integrity of uploaded files

**Supplementary figure 2 Advanced Analysis module in EAP.**

ser can choose the downstream analysis tools implemented in EAP and customize the configure parameters. The input and output examples demonstrate the comprehensive visualization and analysis results.

**Supplementary figure 3 Illustrating the visualization of the TF activity scores of RUNX1/2/3 in two case study ATAC-seq data sets using the Data Set Browser**

## Supplementary movies

**Supplementary_movie_1_Data_transferring_and_integrity_checking.mp 4**

**Supplementary_movie_2_Basic_Analysis_module_demo_using_paired_ end_ChIP_seq_data.mp4**

**Supplementary_movie_3_Advanced_Analysis_module_demo_differentia l_analysis.mp4**

