## Supplementary figures for "EAP: a versatile cloud-based platform for comprehensive and interactive analysis of large-scale ChIP/ATAC-seq data sets"

### Supplementary figure 1

a

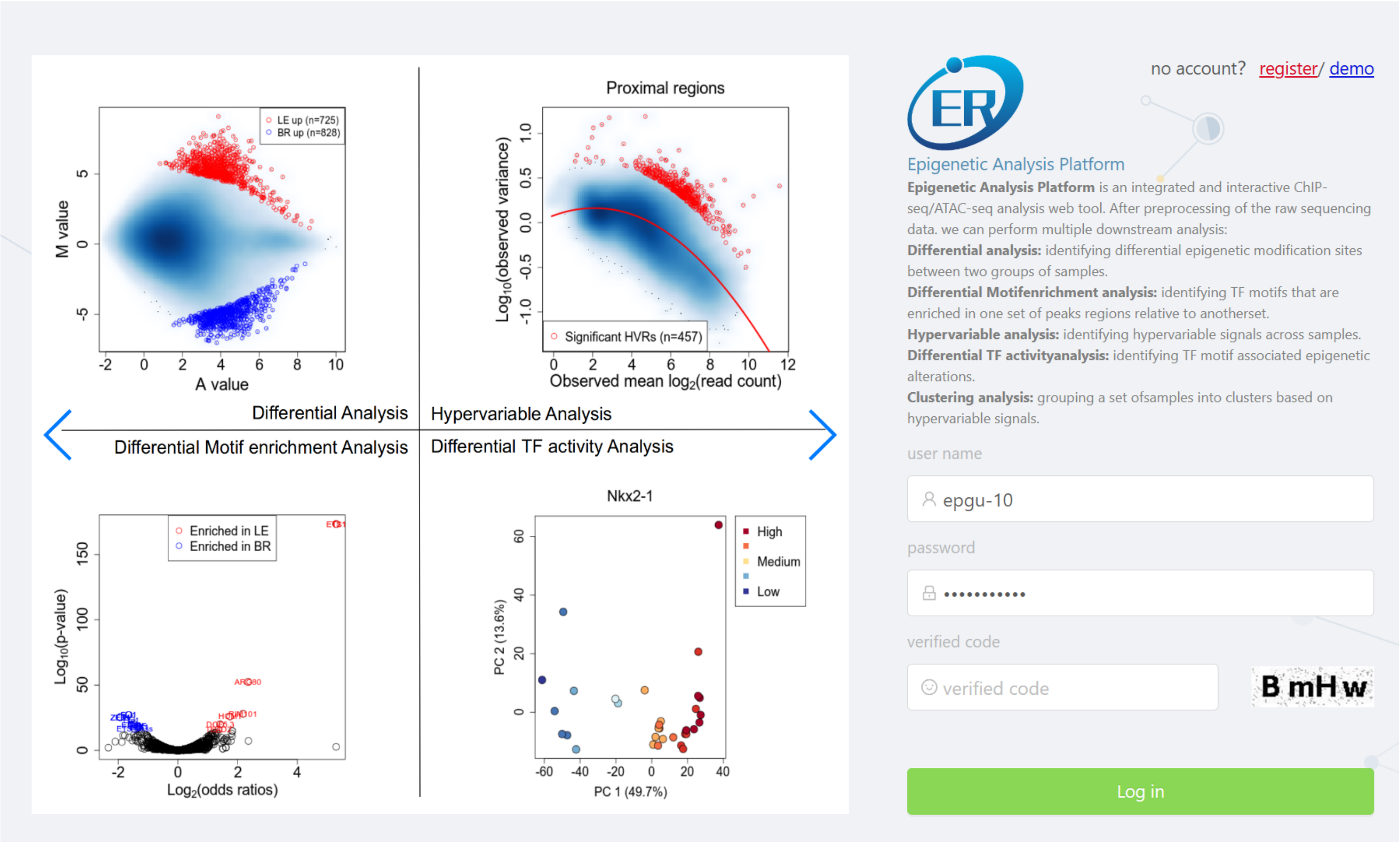


b

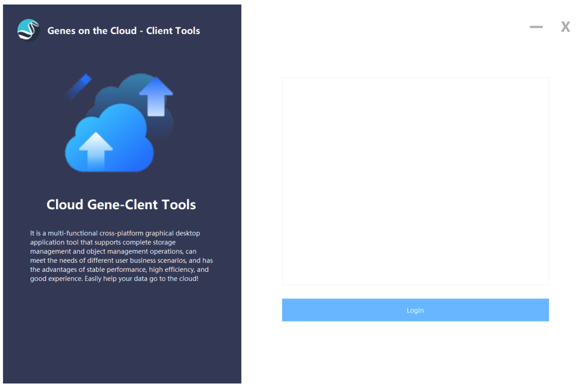


c

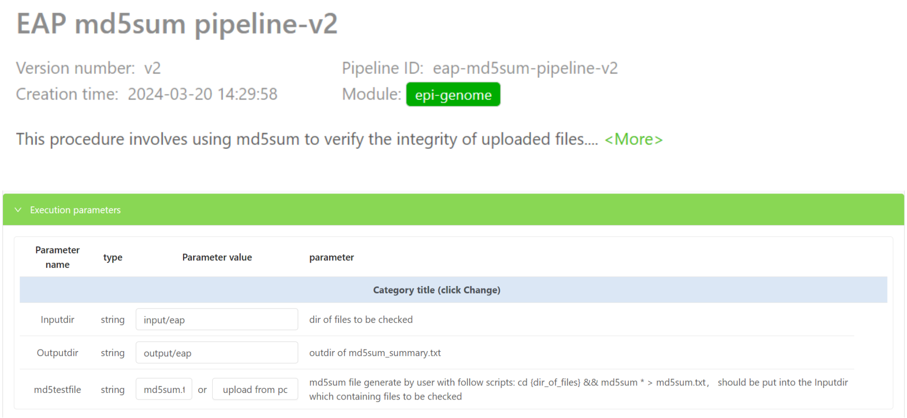

**Supplementary figure 1 The login page and data transfer tools of EAP**

**(a)** The login page of EAP (https://www.biosino.org/epigenetics/#/user/login), a demonstration account is available by click on "demo" in this page. **(b)** The data transfer client for utilization of EAP, which supports break-point resumable transfers. **(c)** A md5sum checking procedure has been integrated to ensure the integrity of uploaded files

### Supplementary figure 2

#### 1) Downstream analysis tools implemented in Advanced Analysis module

**differential-analysis-prd**

Differential analysis: This tool enables the comparison of epigenetic profiles between two distinct conditions and identifies regions that exhibit differential signals.

shaolab | 2023-11-01 21:22:15 | Execute

**diff-tf-enrichment-analysis-prd**

Differential TF motifs enrichment analysis: This tool was used for identifying TF motifs that are enriched in one set of peaks regions relative to another set.

shaolab | 2023-11-02 15:06:43 | Execute

**hypervariable-analysis-prd**

Hypervarable analysis: The Hypervariable Analysis tool detects hypervariable signals in different samples. It conducts the analysis in both proximal and distal peak regions. Principal Component Analysis is then performed based on these identified hypervariable signals. The samples are visualized using a Two-dimensional PCA-scores plot (PC1 vs PC2) or a Two-dimensional TSNE plot, with the color indicating the categorical variable.

shaolab | 2023-11-01 22:48:12 | Execute

**clustering-analysis-prd**

Clustering analysis: grouping a set of samples into clusters based on hypervariable signals. Additionally, a new meta data file will be created that includes cluster labels. With this clustering result, users can conduct differential analysis, differential motif enrichment analysis, and differential TF activity analysis.

shaolab | 2023-11-01 22:57:20 | Execute

**diff-tf-activity-prd**

Differential TF activity analysis: This tool is designed to identify TF motifs that are associated with epigenetic alterations. It achieves this by conducting motif enrichment analysis specifically within hypervariable peak regions. Additionally, it estimates the activity of TFs by aggregating the signals from peaks that are associated with the corresponding TF motif. Finally, it identifies condition-specific TF motifs based on their activity.

shaolab | 2023-11-01 23:08:26 | Execute

**sign-gene-score-prd**

Signature genes score analysis: calculating a signature score from a list of genes for each sample. Signature genes could be downloaded from MSigDB as in GMT format or defined by user based on genes of interest.

shaolab | 2023-11-01 23:10:43 | Execute

#### 2) User-friendly interface for customizable advanced analysis

Parameter setting | IO demo

**clustering-analysis-prd**

Download Template.

Task Information

Inputdir: input/clustering-analysis-prd/1710916704268

Outputdir: output/clustering-analysis-prd/1710916704268

* Proximal peak regions file: input file name or upload from pc

* Distal peak regions file: input file name or upload from pc

* Metadata file: input file name or upload from pc

* Categorical variable: cancer_type

* Name of output clustering result file: Test

* The number of clusters: 2

* Adjusted p-value: 0.1

* The number of PC: 0

Run | Reset

#### 3) Detailed description of input and output from each tool

Parameter setting | IO demo

Input files | Ouput files

In the standard analysis pipeline, this analysis required three input files (proximal and distal peaks were analyzed separately). Two of these input files were derived from the hypervariable analysis, saved in RData format. Each input file contained information on the genomic position of each peak in the first three columns. The remaining columns consisted of a normalized counts matrix, an occupancy matrix, and the result of the hypervariable testing. Users have the option to either upload a local input file or select a file from the server.

| chrom | start | end | sample1.read_cnt | ... | sample1.occupancy | ... | observed.var | prior.var | fold.change | proximal/distal_p_values | proximal/distal_fdrs |
|---|---|---|---|---|---|---|---|---|---|---|---|
| chr1 | 1 | 1000 | 0.2 | ... | 1.3 | ... | 1 | 2 | 2 | 0.001 | 0.01 |
| ... | ... | ... | ... | ... | ... | ... | ... | ... | ... | ... | ... |
| chr22 | 10 | 1000 | 0.5 | ... | 2.5 | ... | 2 | 1 | 0.5 | 0.001 | 0.01 |

The Metadata file was another input file containing information about the samples from the input file. This includes details such as the tissue type of each sample. Users have the option to upload a local input file or select a file from the server.

| Sample_ID | ... | tissue_type | gender | ... |
|---|---|---|---|---|
| sample1 | ... | tumor | male | ... |
| ... | ... | ... | ... | ... |
| sampleN | ... | normal | female | ... |

Parameter setting | IO demo

Input files | Ouput files

1) Figures:

The clustering result based on top-ranked principal components from hypervariable peak regions was visualized using a hierarchical tree plot.

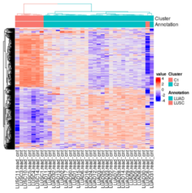


A rank plot displays the optimal number of principal components recommended for hierarchical clustering. However, the user has the option to specify the number of principal components used for clustering.

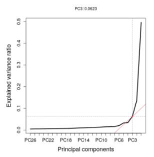


2) Table:

The output file contains the clustering result generated from the clustering analysis. This clustering result has been appended to the original metadata file, creating a new metadata file that can be utilized for other analyses.

| Sample_ID | ... | tissue_type | gender | ... | cluster_ID |
|---|---|---|---|---|---|
| sample1 | ... | tumor | male | ... | 1 |
| ... | ... | ... | ... | ... | ... |
| sampleN | ... | normal | female | ... | 2 |

**Supplementary figure 2 Advanced Analysis module in EAP**

User can choose the downstream analysis tools implemented in EAP and customize the configure parameters. The input and output examples demonstrate the comprehensive visualization and analysis results.

### Supplementary figure 3

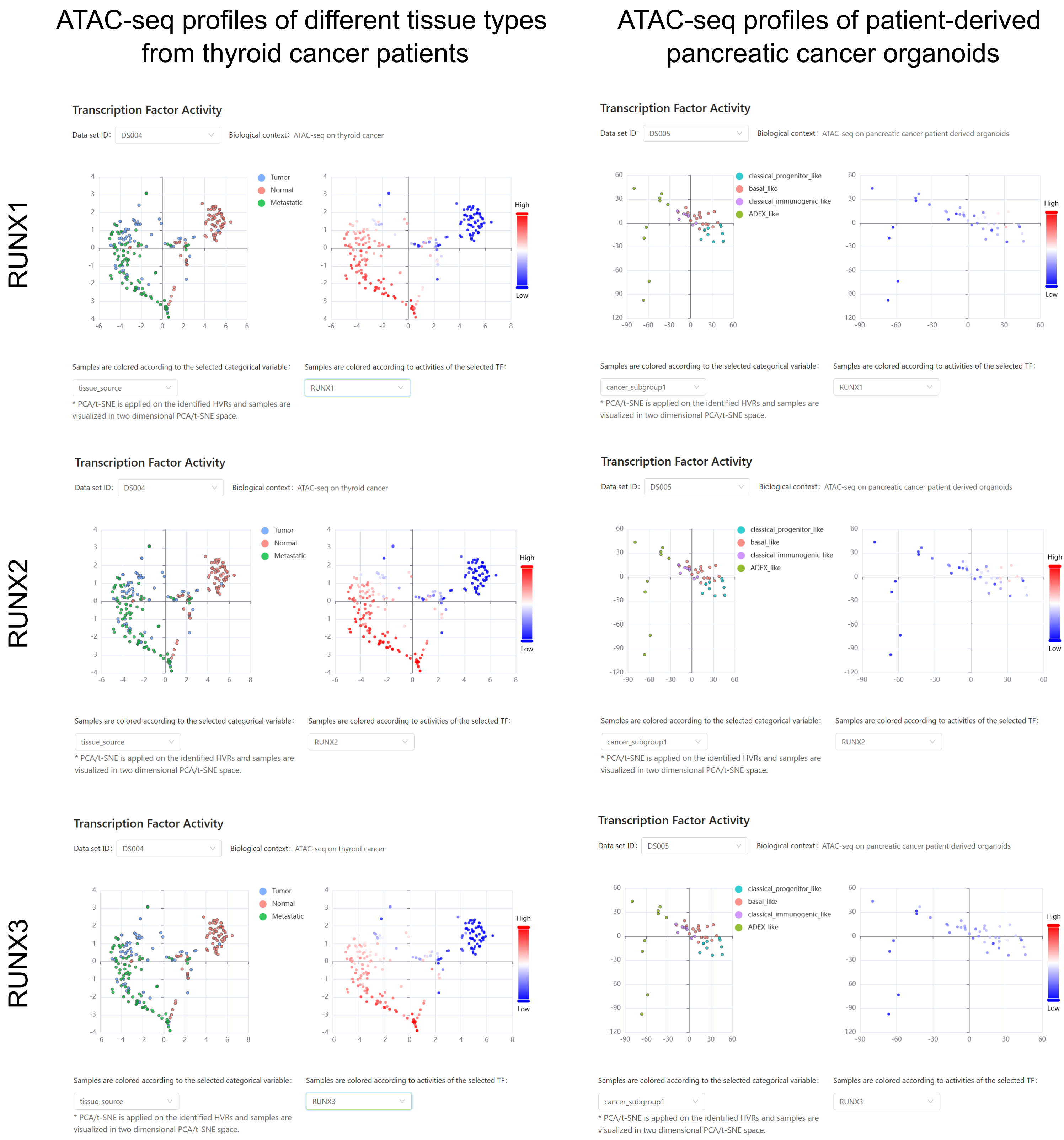

**Supplementary figure 3 Illustrating the visualization of the TF activity scores of RUNX1/2/3 in two case study ATAC-seq data sets using the Data Set Browser**
